# Electrophysiological Signature of Stroke Recovery: Investigating EEG Biomarkers for Prognostic Insights

**DOI:** 10.64898/2026.06.01.728505

**Authors:** Majid Khalili-Ardali, Vivek Sharma, Tarandeep Singh Mandahar, Francisco Páscoa dos Santos, Paul Tiesinga, Nick Ramsey

**Author notes:** Corresponding authors: Paul Tiesinga and Nick Ramsey. Shared first authors (Majid Khalili-Ardali, Vivek Sharma).

## Abstract

Stroke is a leading cause of long-term disability, often resulting in persistent motor impairments reflecting disruptions in large-scale brain networks. While electroencephalography (EEG) has long been used to monitor neurophysiological changes following stroke, an integrated framework capturing spatiotemporal dynamics would help understand changes over time. In this study, we analysed resting-state EEG from stroke patients at one week (Session 1) and three months (Session 2) post-stroke to investigate electrophysiological biomarkers of motor recovery, indexed by changes in the Fugl-Meyer scale (Δ*FM* ). We quantified spectral properties, focusing on the relative alpha band power, microstate metrics such as mean duration, complexity, and transition probabilities, and measures of metastability and synchrony derived from the Kuramoto Order Parameter. Among all the EEG measures, the longitudinal change in relative alpha power emerged as the strongest single correlate of motor improvement, accounting for the largest proportion of variance among the examined EEG measures. Although metastability and synchrony alone did not reach statistical significance, they showed moderate positive correlations with Δ*FM*, particularly in the alpha and theta ranges, and once combined with alpha power, added 26% in explaining the variance in Δ*FM* . Microstate parameters did not explain additional variance in recovery once alpha power and network-level dynamics were considered. A hierarchical model combining alpha power, metastability/synchrony, and microstates explained over 78% of the variance in Δ*FM*, indicating that stroke recovery involves restoring balanced alpha oscillations and flexible large-scale brain coordination. Future research with larger samples and more frequent longitudinal assessments is required to confirm the prognostic utility of integrated EEG biomarkers for guiding personalised stroke rehabilitation strategies.

## 1 Introduction

It is widely recognised that at rest the brain remains highly active, displaying intricate and ever-changing spatiotemporal patterns of activity (Raichle et al. (2001); Fox (2010); Biswas and Banerjee (2025)). This spontaneous neural activity is fundamental to brain function, as it predicts behaviour and reflects intrinsic network dynamics (Cabral et al. (2014)). However, brain injuries such as stroke disrupt these network dynamics, leading to impairments in sensory, motor, and cognitive functions (Salvalaggio et al. (2020)). Motor impairments, in particular, are the most common consequence of stroke, affecting up to 85% of patients in the acute phase and significantly limiting their autonomy in daily activities (Bajaj et al. (2015); Baldassarre et al. (2016); Ramsey et al. (2017)). Understanding how stroke alters large-scale brain dynamics could be of significant value for identifying neural markers of recovery and optimising rehabilitation strategies.

Neurophysiological approaches such as electroencephalography (EEG) provide insights into stroke-related brain dysfunction and altered temporal dynamics of neural activity (Khanna et al. (2015)). Quantitative EEG measures have shown prognostic potential in stroke. For instance, global slowing of EEG rhythms – often indexed by a high delta-to-alpha power ratio – correlates with acute stroke severity and worse outcomes (Sheorajpanday et al. (2011)). Likewise, interhemispheric asymmetry in alpha power (quantified by the Brain Symmetry Index) has been linked to motor impairment, with more asymmetric alpha indicating poorer upper extremity function (Wang et al. (2022)). Beyond traditional spectral and asymmetry measures, more recent analytical approaches have expanded our ability to characterise dynamic brain states. One such well-established method is EEG microstate analysis (Lehmann (1987)), which identifies quasi-stable topographical patterns in scalp electrical activity, which are thought to represent transient global network states (Koenig et al. (2002); Michel and Koenig (2018)). These microstates, which typically last between 40 and 120 milliseconds, are thought to reflect fundamental neural processes underlying cognition and behaviour (Britz et al. (2010); Pascual-Marqui et al. (1995)). While microstate analysis has been extensively applied to psychiatric and neurodegenerative disorders such as schizophrenia and Alzheimer’s disease (Tait et al. (2020); Musaeus et al. (2019)), its potential in understanding stroke recovery has only recently begun to be explored. Early work has pointed out that acute phase microstate patterns might have prognostic value in stroke (Zappasodi et al. (2017)). Additionally, it has been proposed that subacute stroke patients exhibit distinct microstate characteristics in comparison with healthy controls, and the microstate metrics (such as duration, transition probabilities, and complexity) correlate with motor function and recovery (Wang et al. (2022); Hao et al. (2022)).

Beyond spatiotemporal correlates, metastability — the balance between stable and flexible brain states — has emerged as a critical marker of functional integration and segregation in brain networks (Hancock et al. (2025); Deco et al. (2015)). Metastability describes the brain’s ability to transition between transiently stable states while maintaining dynamic flexibility, a feature essential for adaptive behaviour and recovery following brain injury (Kelso (2012); Cabral et al. (2017)). Recent advances suggest that alterations in metastability may provide insights into network reorganisation after stroke and could serve as an indicator of functional recovery (Hancock et al. (2025); Shanahan (2010)). Another key measure of large-scale brain dynamics is synchrony, which refers to the temporal coordination of neural oscillations across different brain regions (Deco et al. (2015)). Synchrony plays a crucial role in motor control and sensorimotor integration, and disruptions in synchrony following stroke have been linked to motor deficits (Trujillo et al. (2017)). In particular, a reduction in interhemispheric alpha coherence has been associated with poorer motor function in daily activities (Kawano et al. (2017)). In contrast, it has been suggested that abnormally high intrahemispheric connectivity may reflect minimal coupling that limits motor recovery (Hoshino et al. (2020)). This growing evidence on the importance of large-scale dynamical features of EEG recordings points to how EEG synchrony measures could indicate how network integration (or its imbalance) contributes to motor impairment and may provide a more comprehensive understanding of stroke-related functional changes and inform rehabilitation strategies.

These findings underscore that EEG can sensitively reflect functional deficits post-stroke. Although previous studies have examined microstate dynamics, metastability, synchrony, and spectral features separately, a unified framework that integrates the large-scale network dynamics with the spatiotemporal pattern of EEG activity and traditional spectral features is still lacking. In this study, we analyse resting-state EEG in stroke patients to investigate changes in microstate dynamics, metastability, synchrony, and classical spectral measures as potential markers of motor recovery. Specifically, we examine how microstate characteristics—such as duration, occurrence, and transition probabilities—are modulated after stroke and whether these changes correlate with functional recovery. Given that microstates reflect transient states of large-scale neural networks, their disruption may provide critical insights into altered brain dynamics post-stroke. Additionally, we assess metastability and synchrony, which capture the temporal stability and coordination of neural oscillations, respectively. While spectral measures are well-established in clinical EEG research, spatiotemporal features (as measured by microstate analysis) and large-scale network dynamics (as measured by metastability and synchrony analysis), despite extensive modeling studies, remain less investigated in clinical applications for stroke populations. Studying these dynamics in clinical settings could yield novel insights into network reorganisation during recovery. Our study aims to characterise each metric independently (microstate parameters, metastability, synchrony, and spectral features) and to develop a unified multivariate model which integrates all these features to better understand the neurophysiological mechanisms underlying stroke rehabilitation. These findings could contribute to the development of more effective therapeutic interventions, ultimately guiding personalised rehabilitation strategies based on neurophysiological biomarkers.

## 2 Methodology

### 2.1 Data Acquisition

A total of 20 stroke patients (mean age 61 years, 9 females) with a first-ever unilateral ischemic stroke in the territory of the middle and/or anterior cerebral artery were enrolled in this cohort. Three patients were excluded (due to missing follow-up scores), leaving 17 patients for final analysis. Electrophysiological and behavioural data were collected at two time points: one week post-stroke onset (Session 1) and three months post-stroke onset (Session 2). Absolute Recovery (AR), as a measure of improvement and functional recovery, was extracted from the normalised Fugl-Meyer (FM) scale (Fugl-Meyer et al. (1975)) and computed as the difference (Session 2 - Session 1) between the normalised FM. Since subjects were largely functionally intact on the ipsilesional side, we only used the Fugl-Meyer motor scores for the contralesional side relative to the lesion.

EEG was recorded using a 128-channel system while participants rested awake with their eyes closed. The dataset used in this study was provided by Adrian Guggisberg and made available through the AISN consortium framework, and was previously published in Dubovik et al. (2012, 2013). The present work did not include new data collection or interaction with human participants. For full details of the original data acquisition, including ethical approval and informed consent, consult Dubovik et al. (2012).

### 2.2 EEG Data Preprocessing

The EEG preprocessing was carried out in Matlab code (R2023a; The MathWorks, Natick, MA, USA) using the Fieldtrip toolbox (Oostenveld et al. (2011)). We applied a zero-phase, two-pass, 4th-order Butterworth IIR filter with a passband from 0.5 to 45 Hz, which was followed by detrending the signal and a notch filter to remove the line noise at 50 Hz. Noisy segments of the data were rejected by visual inspection and using quality metrics (e.g., kurtosis, db amplitude, variance, 1/variance, and z-score), followed by common average referencing (Oostenveld et al. (2011)). Rejected segments never exceeded 10 seconds in length, with total rejection per subject remaining below 1 minute, leaving at least 3 minutes of clean data. All 128 channels were preprocessed, and bad channels were interpolated using nearest neighbour referencing. Approximately 10% of electrodes were labelled as bad channels. We applied *fitting oscillations* and *one over f* (FOOOF) on the power spectral densities (PSD) extracted from the preprocessed EEG data to remove the 1/f power and only analyse the oscillatory part of the signal (Gerster et al. (2022)). All the subsequent analysis steps were performed on this periodic component only. For the frequency band ranges, we used delta (0.5–4 Hz), theta (4–8 Hz), alpha (8–13 Hz), and beta (13–30 Hz) (Cabral et al. (2022)). The relative band powers were calculated as the area under the curve in the specific band range divided by the total power and correlated with the Δ*FM* using Pearson correlation coefficients Oostenveld et al. (2011).

### 2.3 Microstates

We used the +microstate toolbox for calculating the EEG microstates (Tait and Zhang (2022)), which uses a modified k-means clustering method, based on topographical dissimilarity instead of the Euclidean distance. First, the standard deviation of the voltages across all channels at each time point, known as the Global Field Power (GFP), was extracted from the preprocessed EEG data as follows:

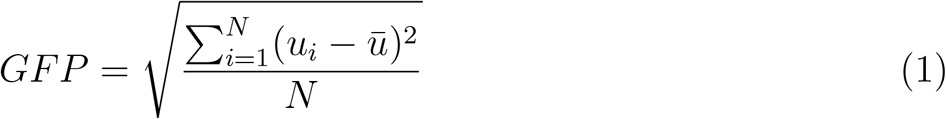

Where *u*_*i*_ is the voltage at electrode *i, ū*_*i*_ is the average voltage of all electrodes, and N is the number of electrodes. Once the GFP was obtained for the entire EEG recording, topographic maps at the local maxima of the GFP in time were clustered via a modified k-means algorithm. We determined the optimal number of clusters to be six using the silhouette score Rousseeuw (1987). Real-time EEG microstates were then derived post hoc by fitting these cluster maps back to each time point in the data, and the statistical significance of the classification accuracy was tested with 100 permutations, for details see (Michel and Koenig (2018)). Because lesion side, location, and recovery severity varied among patients, and because we hypothesise that microstate parameters correlate with network dynamics and recovery, microstates were computed individually for each subject and session. This approach assumes that microstates can change significantly post-stroke and helps account for inter-subject variability. Figure 1 shows a sample of one-second EEG data segmentation into 6 microstates.

**Figure 1.**
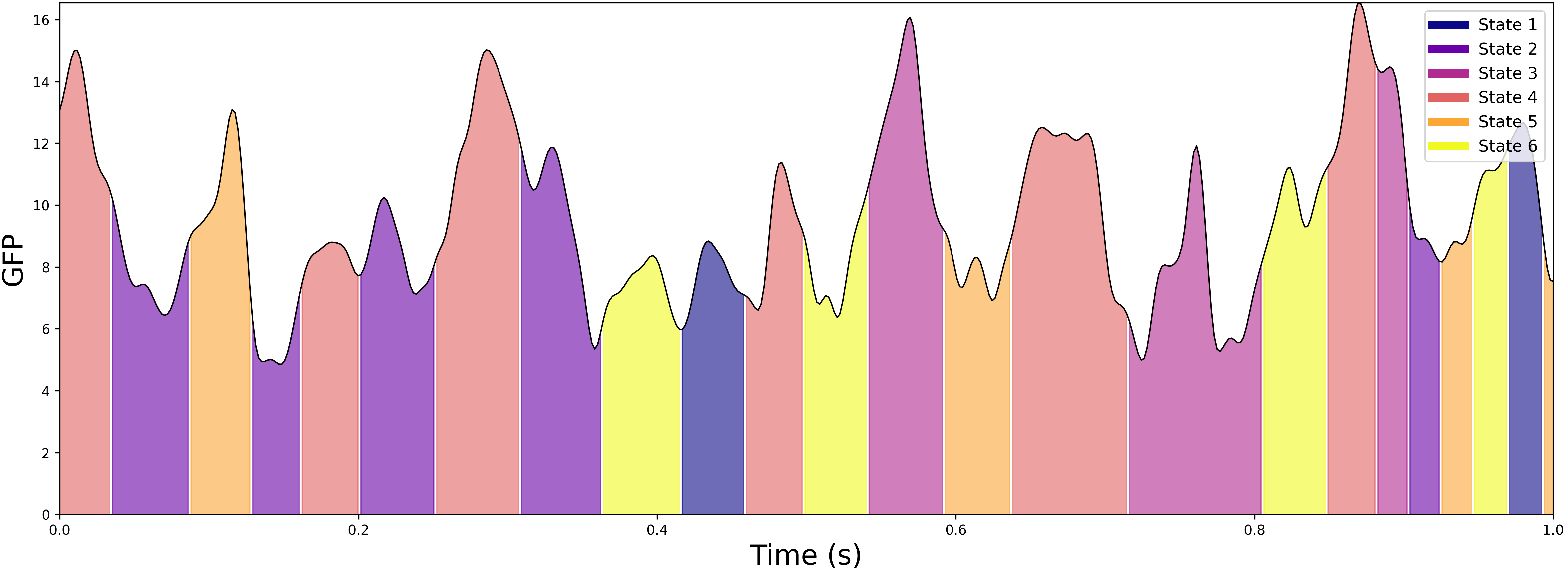
EEG Microstates Segmentation. Global Field Power (GFP) time series overlaid with corresponding microstate segmentation. Each colour-coded segment indicates a distinct microstate cluster derived from spatial topography. Note that GFP reflects the global variance across channels at each time point and does not necessarily distinguish between microstates, which are defined by spatial configuration rather than amplitude differences. Repeated GFP ranges across states reflect distinct spatial patterns occurring at similar global variance levels.

Once the microstate time series was calculated for each subject and session, six measures were derived and correlated separately with the Δ*FM* using Pearson correlation coefficients: (1) the average microstate duration, (2) the Hurst exponent—an index of long-range correlations in time-series data (Van De Ville et al. (2010)), (3) the Lempel–Ziv-inspired complexity—capturing the algorithmic or sequence complexity (Tait et al. (2020)), (4) the GFP peak frequency, (5) the global explained variance (GEV)—which indicates how much of the original EEG signal is explained by the identified microstates (Hu et al. (2022)), and (6) the state transition probability matrix. For the transition probability matrix, the average duration of all microstates (computed separately per subject and session) served as a duration threshold; a transition from the current state to the next (including the same state) was counted whenever the time series remained in one state for at least this threshold. Only transitions with probabilities exceeding those of a baseline model are displayed in Figure 3 and Figure S1 (Supplementary Materials). This baseline model was generated by performing 1000 permutations of the transition matrices for each subject and session, creating a null distribution to assess statistical significance. Once we obtain the null distribution after running these permutations, we proceed to compute the significant differences in the observed transition probability matrix, as compared to the null distribution. For each transition probability *tp*(*i, j*) in the observed 6×6 transition matrix (representing transitions between 6 microstates), we compute the null mean (expected transition probability under the null hypothesis) and standard deviation as:

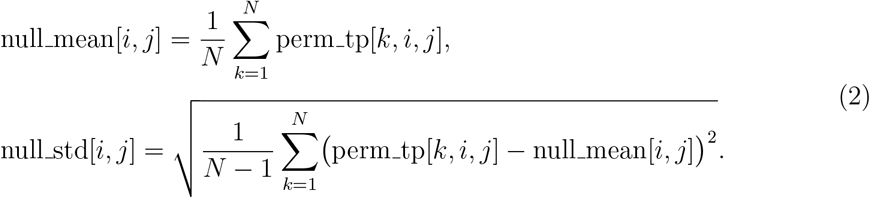

where *N* = 1000 is the number of permutations and perm tp[*k, i, j*] is the transition probability for states *i* to *j* in the *k*-th permutation.

We then calculate the z-score for each transition, assuming the null distribution is approximately normal:

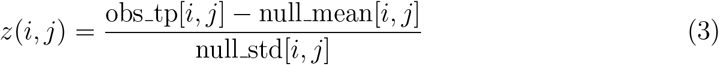

where obs_tp[*i, j*] is the observed transition probability.

To account for multiple comparisons across the 36 transitions in the 6×6 matrix, we apply a Bonferroni correction to control the family-wise error rate. With a nominal significance level of *α* = 0.05, the adjusted threshold is *α*_bonf_ = *α/*36 ≈ 0.00139. Transitions with *p*(*i, j*) *< α*_bonf_ are deemed statistically significant.

### 2.4 Metastability and Synchrony

To analyse oscillatory dynamics, EEG signals were bandpass filtered into four standard frequency bands: delta (0.5–4 Hz), theta (4–8 Hz), alpha (8–13 Hz), and beta (13–30 Hz). The filtering process was applied separately for each trial using a second-order Butterworth bandpass filter implemented with zero-phase forward and reverse filtering to prevent phase distortions (Gustafsson (1996)). Following bandpass filtering, the signals were orthogonalised (Hipp et al. (2012)) to remove shared variance between channels, ensuring an independent phase estimation. The orthogonalised signals were then subjected to a Hilbert transform to extract instantaneous phase information, and the amplitude of the analytic signal was normalised to obtain pure phase values.

Metastability and synchrony were assessed using the Kuramoto Order Parameter (KOP), a well-established measure of global phase synchrony (Hancock et al. (2025)). The KOP at each time point quantifies the degree of phase alignment across electrodes and is computed by averaging the unit-norm complex exponentials of the phase angles (Hancock et al. (2025); Cabral et al. (2022)).

The Kuramoto Order Parameter at time t is given by:

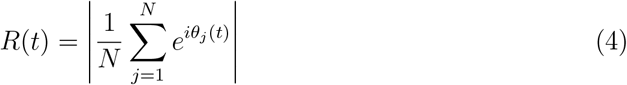

where *θ*_*j*_(*t*) represents the phase of electrode *j* at time *t*, and *N* is the total number of electrodes.

To characterise network dynamics, the mean KOP value over time was used as a measure of global synchronisation, while its standard deviation captured the degree of temporal variability in synchrony, reflecting metastability (Shanahan (2010)).

For each subject and frequency band, the KOP was calculated by computing the phase of each orthogonalised and filtered signal, averaging the unit norm complex exponentials of phase angles across EEG channels, extracting the absolute value of the resulting vector to obtain a time series of phase synchrony values and computing the mean (synchrony) and standard deviation (metastability) of the KOP time series.

To examine the relationship between metastability, synchrony, and motor recovery, the differences in these measures were computed across the two recording sessions. These differences were then correlated with changes in Fugl-Meyer motor scores, a clinical measure of motor recovery. Pearson correlation was used to quantify the strength of the relationship between these variables, computed using the formula:

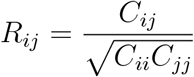

where *C*_*ij*_ represents the covariance between variables *i* and *j*, and *C*_*ii*_ and *C*_*jj*_ are the variances of each variable. Pearson correlation coefficients were extracted from the correlation matrix to evaluate the association between changes in EEG-derived measures and motor recovery outcomes.

### 2.5 Unified Multivariate Framework

We developed a three-block hierarchical regression model to integrate EEG measures from multiple domains (spectral features, microstate metrics, and network-level dynamics) into a single explanatory framework for motor recovery. In each block, we first identified the single best EEG predictor by computing the Pearson correlation between each candidate variable and the Δ*FM* . This filter step ensures that, within each block, only the strongest correlating predictor enters the regression, thus reducing dimensionality in this small-sample dataset. After selecting the top predictor *S* from the spectral block, *M* from the network block, and *N* from the microstate block, we fit a sequence of regression models. First, we regress the outcome Δ*FM* on the chosen spectral predictor *S*:

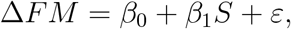

and record the proportion of variance explained 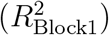 . Next, we add the large-scale oscillatory dynamics predictor *M* to the model:

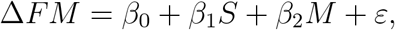

obtaining a new 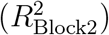. The increment 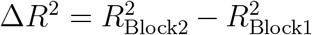 is attributed to large-scale oscillatory dynamics. Finally, we include the microstates predictor *N* in the full model:

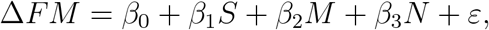

resulting 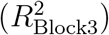 . Each stage’s Δ*R*^2^ quantifies that block’s unique contribution beyond preceding blocks. The final model’s 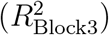 captures how much combined variance in Δ*FM* is explained by the three selected EEG predictors. Additionally, each block’s Δ*R*^2^ indicates the incremental explanatory power contributed by spatiotemporal (microstate) or network (synchrony/metastability) metrics above and beyond the spectral domain. The sequence of block entry was based on an analysis of single-parameter correlations with the outcome variable. While linear regression models are order-independent in terms of final fit and parameter estimation, hierarchical regression is used to assess the incremental variance explained Δ*R*^2^ by different predictor blocks. Therefore, the order in which blocks are entered reflects the theoretical or empirical importance we assign to each domain. In our case, we chose to enter blocks sequentially based on their individual correlation strength with the outcome: starting with spectral features (highest r), followed by network-level dynamics, and finally microstate metrics. This order allows us to quantify how much additional explanatory power each domain contributes beyond the preceding ones. To evaluate potential information overlap, we did not explicitly compute correlations between block summary values, as only the spectral block showed significant single-parameter correlations with the outcome. To further address potential redundancy, we alternated the order of the second and third blocks (network and microstate features) during model fitting, as described in the Results.

This unified framework offers an integrated view of which EEG features—and which domains—account for observed variability in motor recovery. In the spectral domain, we use only relative alpha power as the main feature. In the microstate domain, the five metrics (mean duration, Hurst exponent, global explained variance, complexity, and transition probabilities) were first standardised and then averaged to form a single composite microstate index, which was subsequently used in the hierarchical model. Similarly, in the network-level domain, we standardised and averaged synchrony and metastability (across alpha and delta bands) into one single network index.

## 3 Results

### 3.1 Alpha Peak Shift and Relative Alpha Power

Alpha peak frequency shifts have traditionally been proposed as markers of cortical reorganisation during stroke recovery (Juhász et al. (2009)). In our dataset, however, the maximum alpha peak shift across all channels did not exhibit a significant correlation with motor improvement (r = 0.09, p = 0.71). By contrast, when examining relative alpha band power averaged across channels, showed a robust monotonic association with clinical improvement measured by Δ*FM* (r = 0.72, p = 0.00099). Figure 2 shows the spatial distribution of relative alpha power changes by presenting a topographical scalp map, where each electrode location is colour-coded to reflect the mean difference in relative alpha power (Session 2 minus Session 1) across subjects. We did not observe any correlation between the lesion side and the maximum relative alpha power shift. These findings suggest that while alpha frequency shifts alone may not reliably track functional gains, the proportion of power in the alpha band can provide a more sensitive index of neurophysiological changes linked to stroke rehabilitation.

**Figure 2.**
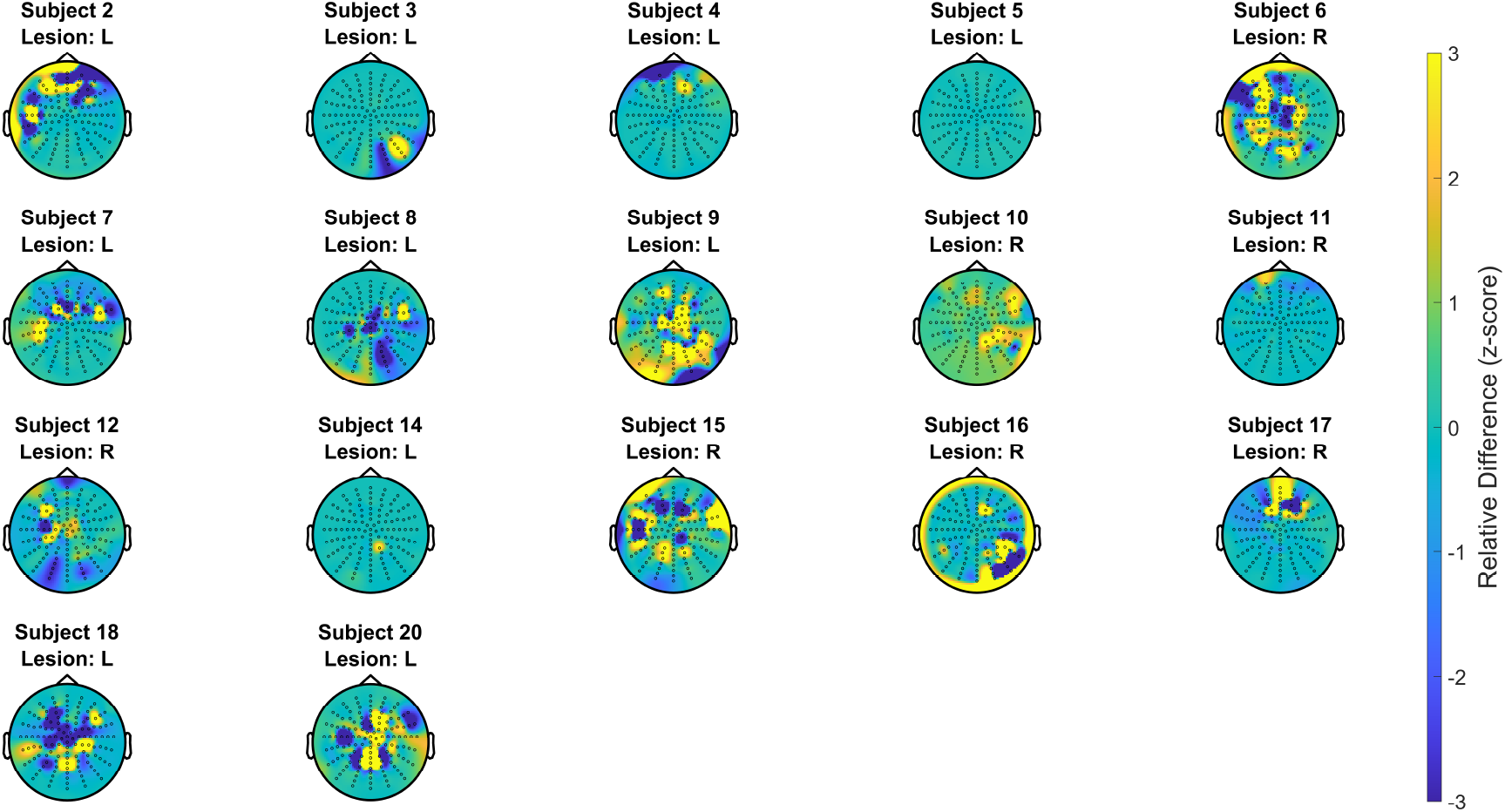
Spatial Distribution of Relative Alpha Power Change. Topographic maps showing the change in relative alpha power from Session 2 to Session 1 for each individual subject. For each subject, relative alpha power was computed at each electrode as the ratio of alpha power to total power, and the difference between sessions (Session 2 – Session 1) is visualised. Each map reflects the spatial pattern of alpha power modulation over time, with warmer colours indicating increases and cooler colours indicating decreases.

### 3.2 Metastability and Synchrony

We examined the relationship between metastability, synchrony, and Δ*FM* scores across different frequency bands. In the delta band, metastability (r = 0.350) and synchrony (r = 0.315) showed moderate positive correlations with Δ*FM* score differences. Similar trends were observed in the theta band, where metastability (r = 0.356) and synchrony (r = 0.418) exhibited slightly stronger associations with Δ*FM* . The alpha band demonstrated the highest correlations, with metastability (r = 0.438) and synchrony (r = 0.417), suggesting that dynamic coordination in this frequency range may be more relevant to recovery. In contrast, the beta band showed weaker correlations, with metastability (r = 0.211) and synchrony (r = 0.246). While none of the correlations reached statistical significance at *α* = 0.05, the observed trends suggest increased metastability and synchrony in lower frequency bands.

### 3.3 Microstate Analysis

All statistical measures extracted from the microstate time series were correlated with the Δ*FM*, and none of them individually reached significance at *α* = 0.05. Mean microstate duration (*r* = 0.09), Complexity (*r* = 0.05), GFP peaks (*r* = −0.18), and GEV (*r* = −0.15) all showed weak correlation relationships with Δ*FM*, suggesting that no single measure from microstates can be used to predict motor recovery solely. Since patients exhibited heterogeneous lesion locations, affected hemispheres, and recovery trajectories, the cortical reorganisation was expected to vary greatly across individuals; hence, we performed subject-specific analyses of transition probabilities between different microstates. As shown in Figure S1 (Supplementary Materials), there seem to be subject-specific changes in microstate transition probabilities between the sessions. Figure 3 illustrates a representative subject whose transition probabilities between Session 1 and Session 2 differed substantially. It should be noted that, in these figures, only transitions exceeding the baseline model are plotted; therefore, more arrows indicate more structured state-to-state transitions. To quantify these results, we computed the average graph density, which increased from 0.65 in Session 1 to 0.70 in Session 2. However, across subjects, the average graph density did not significantly correlate with Δ*FM* (*r* = 0.079, *p* = 0.76). The graph density is computed using the following equation:

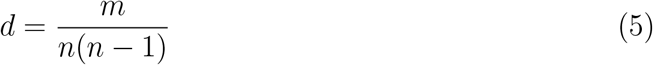

where *n* is the number of nodes and *m* is the number of edges in *G*. The number of connections in the graph is the same as the number of significant transition probabilities.

**Figure 3.**
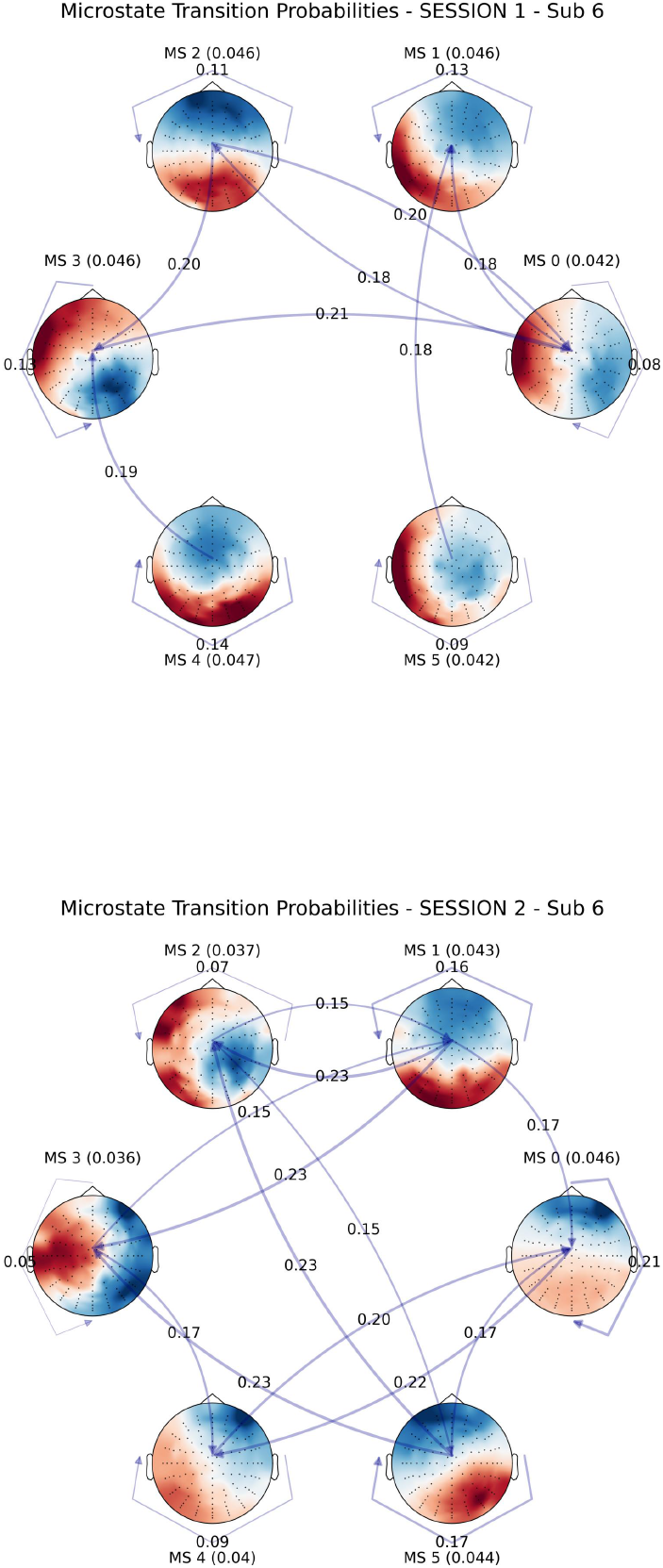
Transition Probabilities for subject 6. Directed graphs showing microstate transition probabilities for Session 1 and Session 2. Microstates were independently extracted for each session, reflecting the expected variability due to lesion effects and recovery over time. As such, microstate labels (MS1–MS6) may not correspond directly between sessions. The number of states was kept constant at six per session, based on silhouette distance optimisation, to enable comparable transition structures. Only transitions that were significantly above chance level, determined via bootstrap resampling, are shown.

### 3.4 Unified Multivariate Framework

In the first-level regression, averaged relative alpha power changes alone accounted for 52% of the variance in motor recovery in a linear regression model (R^2^ = 0.5257, p = 0.0010), indicating a strong association in this sample between these spectral changes and functional outcomes. In the second step, adding the large-scale oscillatory dynamics increased the total explained variance by 26% (*R*^2^ = 0.7861, *p* = 0.0000), suggesting that global dynamics contributed additional information beyond alpha power alone. In the third-level model, incorporating the microstate predictor only improved the explained variance by 0.12% (*R*^2^ = 0.7873, *p* = 0.0001), showing that microstates did not have any additional explanatory value beyond what is captured by metastability and synchrony. In this dataset, microstate features accounted for 3.6% of explained variance when entered as the second block, compared to 26% from large-scale network dynamics (synchrony/metastability). When microstates were entered as the third block, they contributed only an additional 0.12%, suggesting that most of their variance was shared with earlier blocks. While alpha power emerged as the strongest single predictor, the increase in R^2^ across levels highlights the incremental contribution of large-scale oscillatory dynamics.

## 4 Discussion

In this study, we examined how EEG metrics of global dynamical properties (as measured by metastability and synchrony), spatiotemporal features (as measured by microstate analysis), and classical spectral measures (particularly alpha band power) relate to post-stroke changes in Fugl-Meyer scores (Δ*FM* ). Of these, only the relative alpha band power showed a consistent positive association with (Δ*FM* ). Metastability and synchrony demonstrated moderate positive correlations, especially in the alpha and theta bands, suggesting that enhanced large-scale oscillatory coordination may favour better recovery outcomes. In this study, only spectral and network-level features demonstrated clear associations with recovery outcomes. Microstate measures, including duration, global explained variance (GEV), complexity, GFP peak frequency, and transition probability measures, showed low or inconsistent correlations and did not contribute meaningfully to explained variance. Thus, our data do not support strong conclusions regarding the role of microstate dynamics in stroke recovery. Overall, these findings suggest that stroke recovery primarily involves reestablishing functional balance at the spectral and network levels of brain organisation. In our dataset, improvements in recovery outcomes were associated with longitudinal increases in spectral power in the alpha range and elevated measures of network metastability and synchrony. These findings suggest that elevations in spectral and network-level dynamics may reflect ongoing reorganisation processes during motor recovery. We did not observe statistically meaningful contributions of microstate dynamics in this cohort. Overall, these exploratory observations indicate that stroke recovery may be accompanied by enhanced spectral and network properties, though stronger evidence would be required to establish causal relationships or normative thresholds.

Our finding of a positive correlation between relative alpha power increases and motor recovery appears to conflict with reports that suggest higher alpha at single time points portends worse function (Leonardi et al. (2022); Martino Cinnera et al. (2025); Sebastián-Romagosa et al. (2020); Hoshino et al. (2020)). Importantly, our metric captured longitudinal change in alpha. This likely reflects a normalisation of cortical rhythms over time, rather than the static excess alpha seen in acute stroke (‘idle’ cortex) that correlates with impairment. In fact, recent longitudinal studies support this interpretation: patients who show a shift from slow-wave activity toward alpha-beta activity during rehabilitation tend to achieve greater recovery (Trujillo et al. (2017)), in other words, the reduction of EEG slowing (quantified by a flattening of the spectral exponent) was significantly correlated with improved motor function (Lanzone et al. (2022)). Thus, increases in alpha power over the subacute period may signify the restoration of more normal network excitability and engagement, rather than pathological inhibition (Sebastián-Romagosa et al. (2020)). Future research with larger cohorts should clarify the precise role of alpha-band dynamics in post-stroke motor recovery and whether any early EEG spectral indicators – e.g. alpha laterality indices or delta/alpha ratios – have true prognostic value when measured in the acute phase of the stroke.

It has been reported that stroke patients with lower motor function showed prolonged microstates of types A and B compared to controls, while those microstate parameters tended to normalise as function improved (Hao et al. (2022)). Similarly, it has been reported that certain microstate metrics (e.g. fractional coverage of specific classes) were significantly correlated. In our analyses, microstate features showed limited unique contribution once spectral power and large-scale network dynamics were considered. This suggests that much of the variance captured by microstate metrics overlaps with broader dynamical measures, whereas alpha power and network-level synchrony/metastability emerged as the main independent contributors to model performance.

Neurophysiologically, increased metastability and synchrony were correlated with recovery outcomes in our dataset, potentially reflecting a greater degree of freedom for state transitions, indicating that the system state space is more accessible. While the functional significance of these associations cannot be directly discussed from these analyses, it is possible that elevated metastability supports the exploration of a broader repertoire of network configurations, which may in turn facilitate neuroplastic adaptation and functional reorganisation in motor-related circuits following stroke. In parallel, shorter microstate durations and reduced GEV may indicate greater flexibility in neural network organisation, suggesting that the brain is not constrained to a few dominant patterns but can dynamically reorganise in response to rehabilitation efforts. Although metastability has been conceptually linked to measures such as the Kuramoto order parameter, no existing studies have directly examined the relationship between the Kuramoto parameter and Fugl-Meyer outcomes. Future work can explore the Kuramoto order parameter to better understand how oscillatory coupling supports motor recovery. Our results align with previous findings identifying large-scale neural coordination as essential for stroke rehabilitation. For instance, Kawano et al. (2017) and Riahi et al. (2020) demonstrated that increased interhemispheric synchrony or interconnectivity corresponds to reduced motor impairment, supporting our finding that alpha- and theta-band synchrony is associated with improvements in motor recovery. In the Unified Multivariate Framework, relative alpha power changes emerged as the single strongest predictor of motor recovery, explaining 52.57% of the variance; however, adding global network-level features and spatiotemporal microstate metrics incrementally raised the explained variance first to 78.61% and then to 78.73%. This pattern indicates that these different EEG domain features capture distinct aspects of post-stroke cortical reorganisation. While alpha power alone has frequently been highlighted in the literature, our results indicate that including large-scale network coordination refines the explanatory model further, and microstate dynamics do not explain beyond the combination of the first two measures. It should be noted that since the sample size was limited in this study, a composite index for each domain was created by averaging standardised microstate metrics or network-level measures to overcome overfitting issues. Although this domain-level aggregation potentially sacrifices some granularity, it preserves statistical stability in a small-sample setting. Future studies with larger cohorts could more fully exploit the diversity of microstate and network features, offering deeper insight into the mechanisms of stroke recovery.

Some prior studies have reported EEG features with modest prognostic utility. Connectivity measures at four weeks were shown to predict motor scores at eight weeks (Hoshino et al. (2020); Riahi et al. (2020); Lee et al. (2022)), and certain microstate parameters measured acutely were found to relate to recovery at 5 months (Zappasodi et al. (2017)). Based on these observations and the current findings, it is suggested that future large-scale or longitudinal studies systematically assess the prognostic value of integrated EEG biomarkers using both large-scale EEG network dynamics and spatiotemporal patterns combined with traditional spectral features, particularly in the acute–subacute transition. Solely based on the current results, it remains unclear whether these measures simply reflect epiphenomenal changes or play a causal role in enabling motor recovery.

Despite these promising findings, several limitations should be acknowledged. A primary limitation is the small sample size, which may significantly limit the statistical power and the generalisability of our results. Future studies with larger cohorts are necessary to confirm the robustness and reproducibility of these correlations. It is also possible that the absence of strong associations with microstate features reflects underlying heterogeneity in lesion location or stroke subtype; future work incorporating lesion-informed subgrouping or mixed-effects models may help clarify these relationships.

## 5 Conclusion

In conclusion, our findings suggest that in this cohort, longitudinal increases in alpha power, along with enhanced metastability and synchrony, were associated with better post-stroke motor recovery. Microstate features provided minimal independent predictive value in this cohort. Together with prior research, these results support the idea that adaptive neural coordination and spectral dynamics play important roles in regaining motor function. The unified model accounted for a substantial proportion of variability in post-stroke motor recovery, underscoring its potential as an integrative biomarker framework for stroke rehabilitation. Future research should explore whether these EEG-derived measures, individually or in combination, can serve as reliable biomarkers to predict stroke rehabilitation outcomes and guide targeted therapeutic interventions.

## 6 Acknowledgements

This work is supported by the European Union through AISN (Horizon Europe, 101057655) and eBRAINS-Health (Horizon Europe, 101058516). We thank Adrian Guggisberg for generously providing the EEG dataset used in this study and valuable feedback.

## 7 Supplementary Material

**Figure S1.**
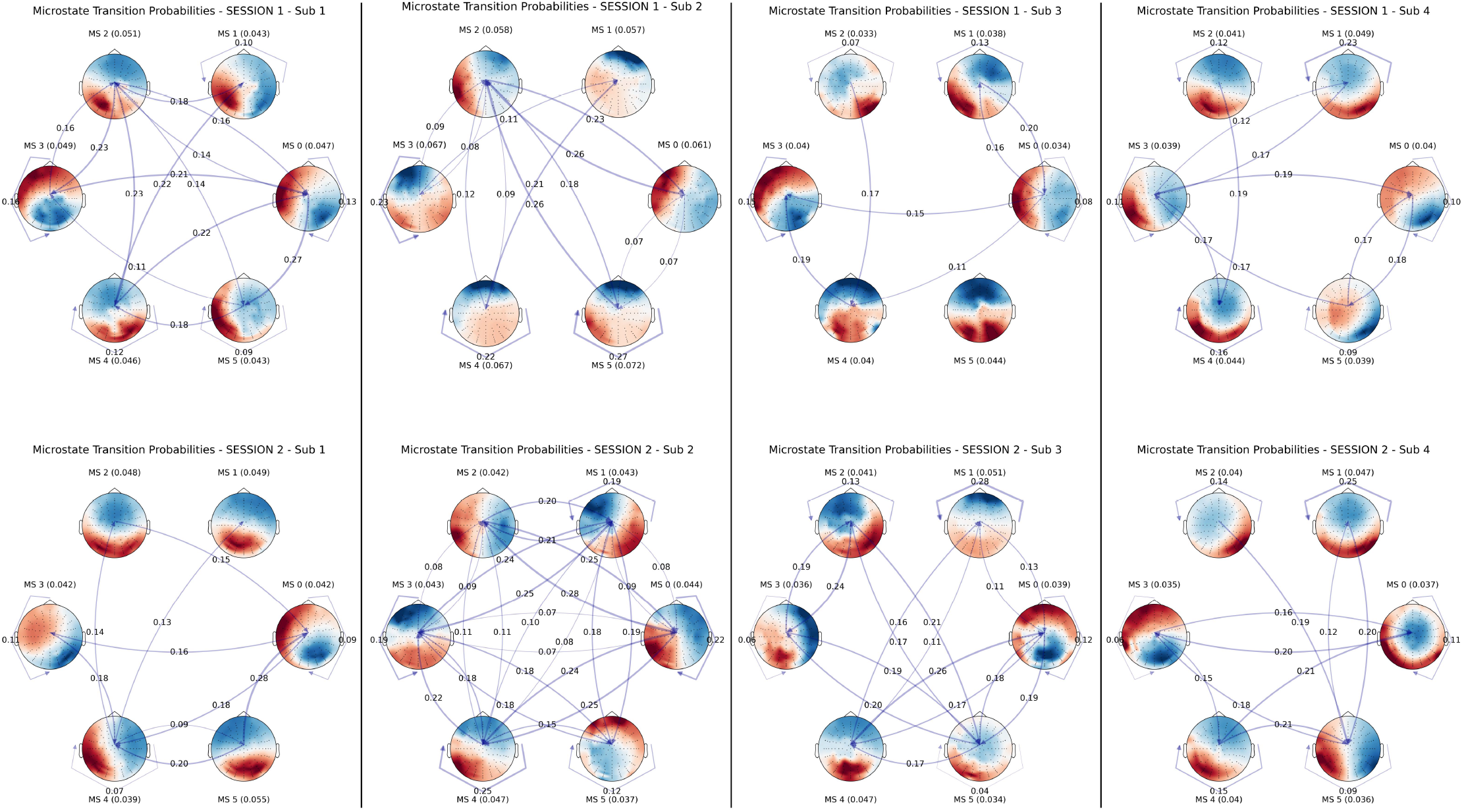
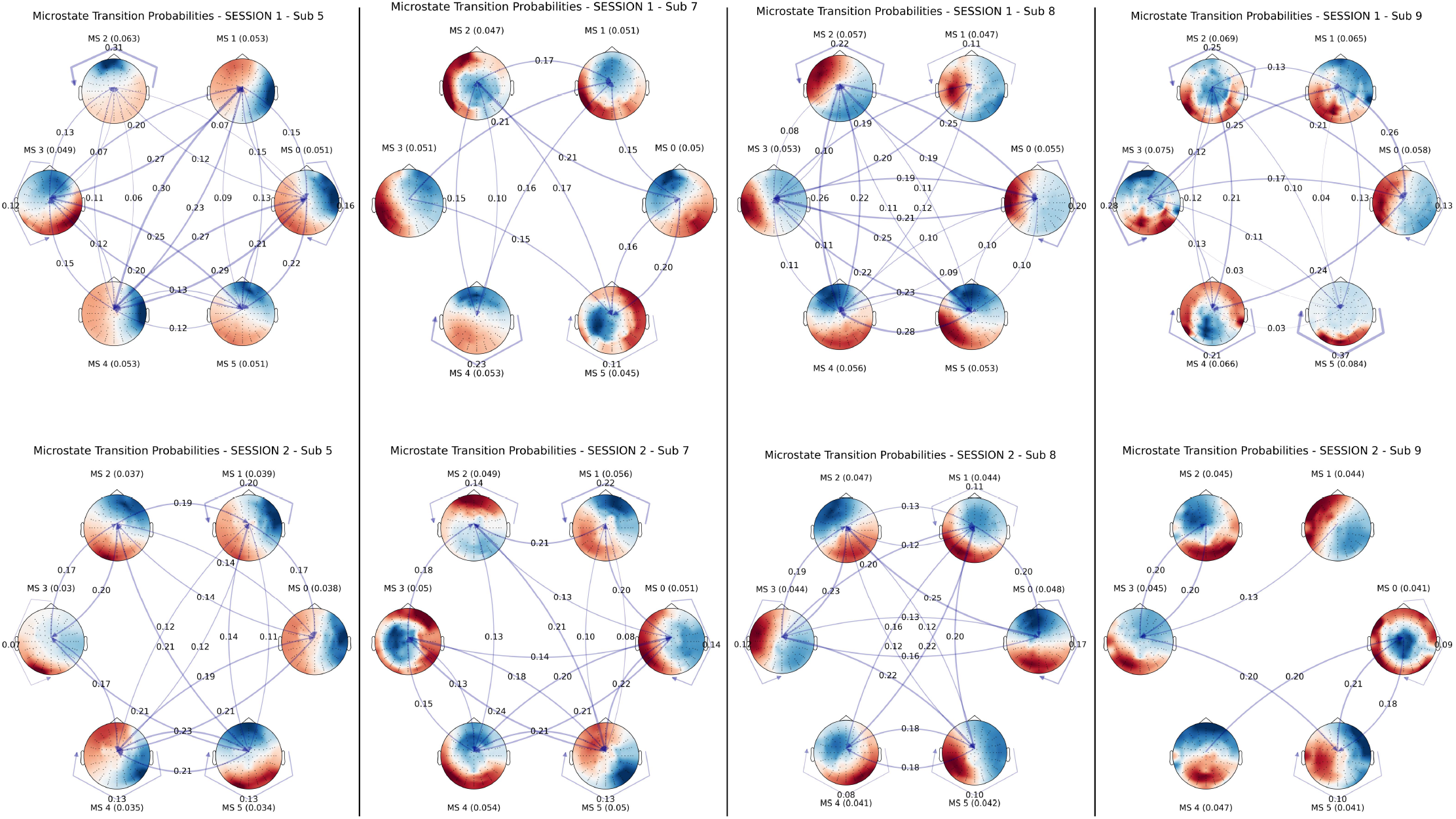
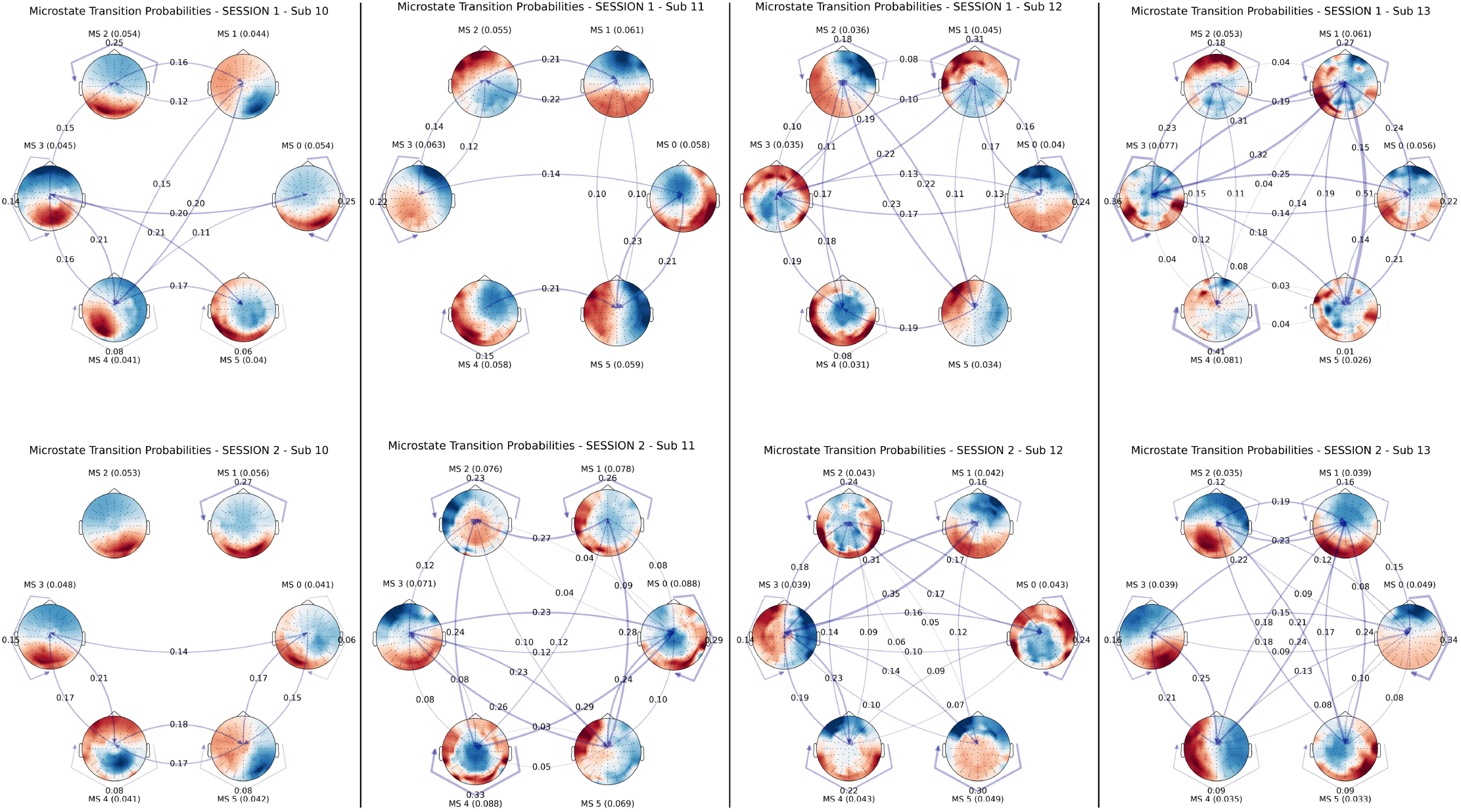
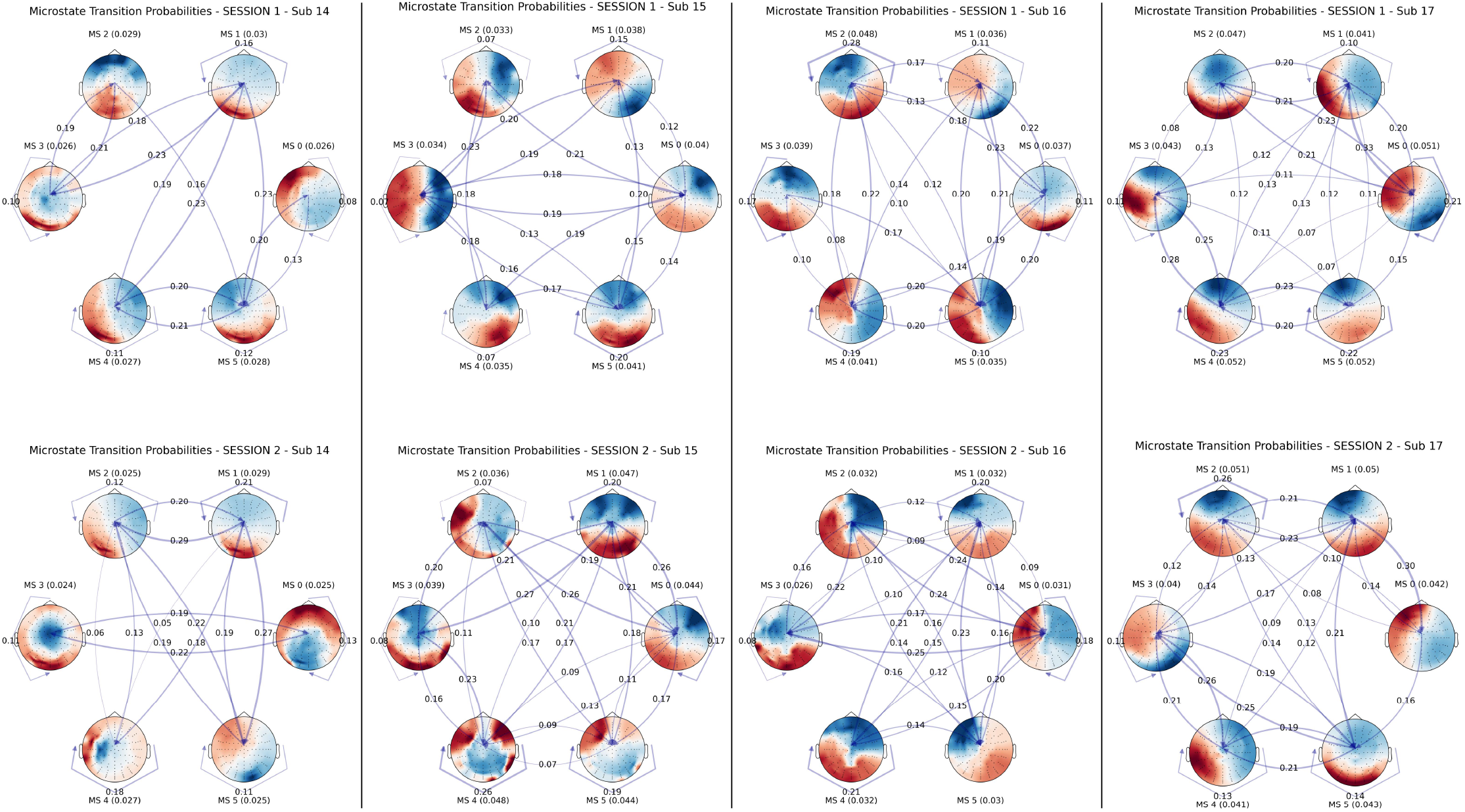
Transition Probabilities for all subjects (columns) and sessions (rows)

